# The luteinizing hormone receptor knock-out mouse as a tool to probe the *in vivo* actions of gonadotropic hormones/receptors in females

**DOI:** 10.1101/2020.08.11.244467

**Authors:** Kim Carol Jonas, Adolfo Rivero Müller, Olayiwola Oduwole, Hellevi Peltoketo, Ilpo Huhtaniemi

## Abstract

Mouse models with altered gonadotropin functions have provided invaluable insight into the functions of these hormones/receptors. Here we describe the repurposing of the infertile and hypogonadal LHR knockout mouse model (LuRKO), to address outstanding questions in reproductive physiology. Using crossbreeding strategies and physiological and histological analyses, we first addressed the physiological relevance of forced LHR homomerization in female mice using BAC expression of two mutant LHR, that have previously shown to undergo functional complementation and rescue the hypogonadal phenotype of male LuRKO mice. In female LuRKO mice, co-expression of signal and binding deficient LHR mutants failed to rescue the hypogonadal and anovulatory phenotype. This was apparently due to the low-level expression of the two mutant LHR and potential lack of LH/LHR-dependent pleiotropic signaling that has previously been shown to be essential for ovulation. Next, we utilized a mouse model overexpressing human chorionic gonadotropin (hCG) with increased circulating ‘LH/hCG’-like bioactivity to ~40 fold higher than WT females, to determine if high circulating hCG could reveal putative LHR-independent actions. No effects were found, thus, suggesting that LH/hCG mediate their gonadal and non-gonadal effects solely via LHR. Finally, targeted expression of a constitutively active FSHR progressed antral follicles to pre-ovulatory follicles and displayed phenotypic markers of enhanced estrogenic activity but failed to induce ovulation in LuRKO mice. This study highlights the critical importance and precise control of LHR and FSHR for mediating ovarian functions and of the potential re-purposing existing genetically modified mouse models in answering outstanding physiological questions.

## INTRODUCTION

The coordinated actions of the gonadotropic hormones, luteinizing hormone (LH) and follicle-stimulating hormone (FSH) and their cognate receptors are essential for reproduction^1,2^. Rare naturally occurring mutations in humans, and laboratory-generated mutations in genetically modified mice, as well as in vitro analyses, have provided vital information on structure-function pairing for gonadotropin hormone/receptor activation, trafficking and signaling^3–7^. However, *in vivo* studies from animal models have proved to be powerful tools for studying the intricacies of gonadotropic hormone/receptor function in a physiologically relevant manner, elucidating their roles in the female in follicle recruitment, selection and growth, ovulation, and corpus luteum function^8–11^. As such, these genetically modified animal models have revealed many important nuances in the molecular and physiological control of reproduction.

A key genetically modified mouse model that has provided important insights into the physiological roles of LHR, is the LHR knockout (LuRKO) mouse^10^. LuRKO animals of both sexes present phenotypically with pubertal delay, hypogonadism and infertility, and have revealed the roles of LHR in pubertal attainment and maintenance of fertility in adulthood^10^. Furthermore, LHR function was found redundant for the prenatal sexual maturation of both sexes^10^. Studies, by us and others, have challenged the central dogmas in the role of LHR in males and females. We therefore aimed to re-purpose and utilize existing previously characterized mouse models of gonadotropin/receptor modifications to address several outstanding questions surrounding the molecular mechanisms by which the LHR and its ligands mediate their physiological functions in females. By utilizing the LuRKO mouse as a background phenotype to be crossed with 3 additional genetically modified mouse models, we first show that female BAC transgenic expression of transactivating mutant LHRs, previously shown to undergo intermolecular cooperation *in vitro and in vivo*^12–14^, failed to alter the infertile female LuRKO phenotype. Second, that transgenic over-expression of the LHR ligand, human chorionic gonadotropin (hCG)^15^, had no LHR independent effects on the hypogonadal phenotype in LuRKO females. Third, expression of constitutively activating mutant (CAM) FSHR^16^ was capable to increase the ovarian follicle development in absence of LHR but failed to induce ovulation and rescue fertility in females, contrasting with male LuRKO mice expressing CAM FSHR^17^.

## METHODS

### Animals

LuRKO mice were produced by targeted disruption of exon 11 of *Lhr* as previously described^10^. Transgenic mice expressing either signal-(LHR^S−^) or binding-deficient (LHR^B−^) form of the Lhr, generated by inserting mutated bacterial artificial chromosomes (BACs), were crossed with LuRKO heterozygotes for 2 generations to obtain LuRKO−/− carrying either one or both transgenes, as described previously^12^. hCGβ transgenic mice were generated using the *ubiquitin C* promoter to drive ubiquitous expression of *hCGβ* transgene as previously described^15^. To generate the double transgenic line with hCGβ expression in a LuRKO background, heterozygous LuRKO animals were inter-crossed with hCGβ expressing mice. The resulting hCGβ/heterozygous LuRKO^−/+^ mice were subsequently inter-crossed to produce hCGβ/LuRKO double transgenic lines. The CAM-FSHR mice were generated by expressing *Fshr*-D580H under the human *anti-Müllerian hormone* promoter^16^. The double mutant Fshr-CAM mice on a homozygous *Lhr*^−/−^ background (LuRKO/CAM FSHR) were generated using a three-step breeding program as previously described^17^. Briefly, the Lhr^+/−^ females were first backcrossed into FVB/N background, and the females produced were intercrossed with the male transgenic Fshr-CAM mice to obtain the Fshr-CAM/Lhr^+/−^ males, which were finally crossbred with the Lhr^+/−^ females in FVB/N background. In this context WT mice referred to mice that after the breeding program did not contain the *Fshr*-D580H transgene and expressed *Lhr*. In the analyses Lhr^+/−^ mice, showing no phenotype deviant from WT, have been pooled together with WT mice and likewise Fshr-CAM/Lhr^+/−^ mice with Fshr-CAM/Lhr^+/+^ mice.

All procedures were carried out in accordance to the regulations of the UK Home Office Animals (Scientific Procedures) Act, the Imperial College London guidelines for animal care, and University of Turku Ethical Committee on Use and Care of Animals approval. To ensure transgene transmission/deletion, ear notches were obtained and genotyped as previously described^10,12,15–17^. To assess for pubertal onset, vaginal opening was monitored daily from day 21 until detected, as previously described^16^. Vaginal smears were investigated daily for 14 days to monitor regularity of estrus cycles.

### Histological analyses

Ovaries and uteri were dissected, weighed and visualized for changes prior to fixation in 4% paraformaldehyde for 4-24 hours depending on size, or snap frozen in liquid nitrogen for gene expression analysis. Fixed tissues were dehydrated in graded ethanol solutions until absolute water-free, cleared in histoclear (National Diagnostics, Hessle Hull, UK) and embedded in paraffin. Tissues were sectioned at 5-μm thickness, mounted on polylysine slides (VWR, Lutterworth, UK), dried at 37 °C for approximately 1 hour, and stored for subsequent use. For histological analysis, tissue sections were stained with the standard hematoxylin and eosin protocol. The estrous cycle was monitored by daily morning analysis of vaginal smears collected for 2-3 weeks, in 8-week-old animals. Vaginal smears were stained by the Giemsa method, and stages defined as proestrus, estrus, metestrus 1–2, and diestrus. All histological samples were imaged using a Nikon Eclipse ME600 with a mounted Nikon D1500 digital camera.

### Quantitative Real-Time PCR

Total mRNA was extracted from ovaries and purified using TRIsure and phenol-chloroform following clean-up with RNeasy kit (Qiagen), including DNase treatment. The purity and quantity of isolated RNA was estimated spectrophotometrically with the use of Nanodrop (ThermoFisher). Reverse transcription (RT) was performed with AMV-reverse transcriptase (Promega). Quantitative PCR (qPCR) reactions were performed using DyNAmo SYBR Green (Finnzymes) kit. PCR reactions were performed in triplicate in a qPCR thermocycler (Chromo4 with OpticonMonitor software, Bio-Rad), using specific primers to LHRs (WT, LHR^B−^ or LHR^S−^).

Sample normalization was assessed vs a housekeeping gene (*Ppia*). A linear standard curve was drawn using different dilutions of a plasmid containing the cDNA of the *Lhr*. Results expressed after adjusting to the housekeeping gene expression: *Lhr*/*Ppia*. Primers for qRT-PCR are shown in Table 1.

**Table 1:**
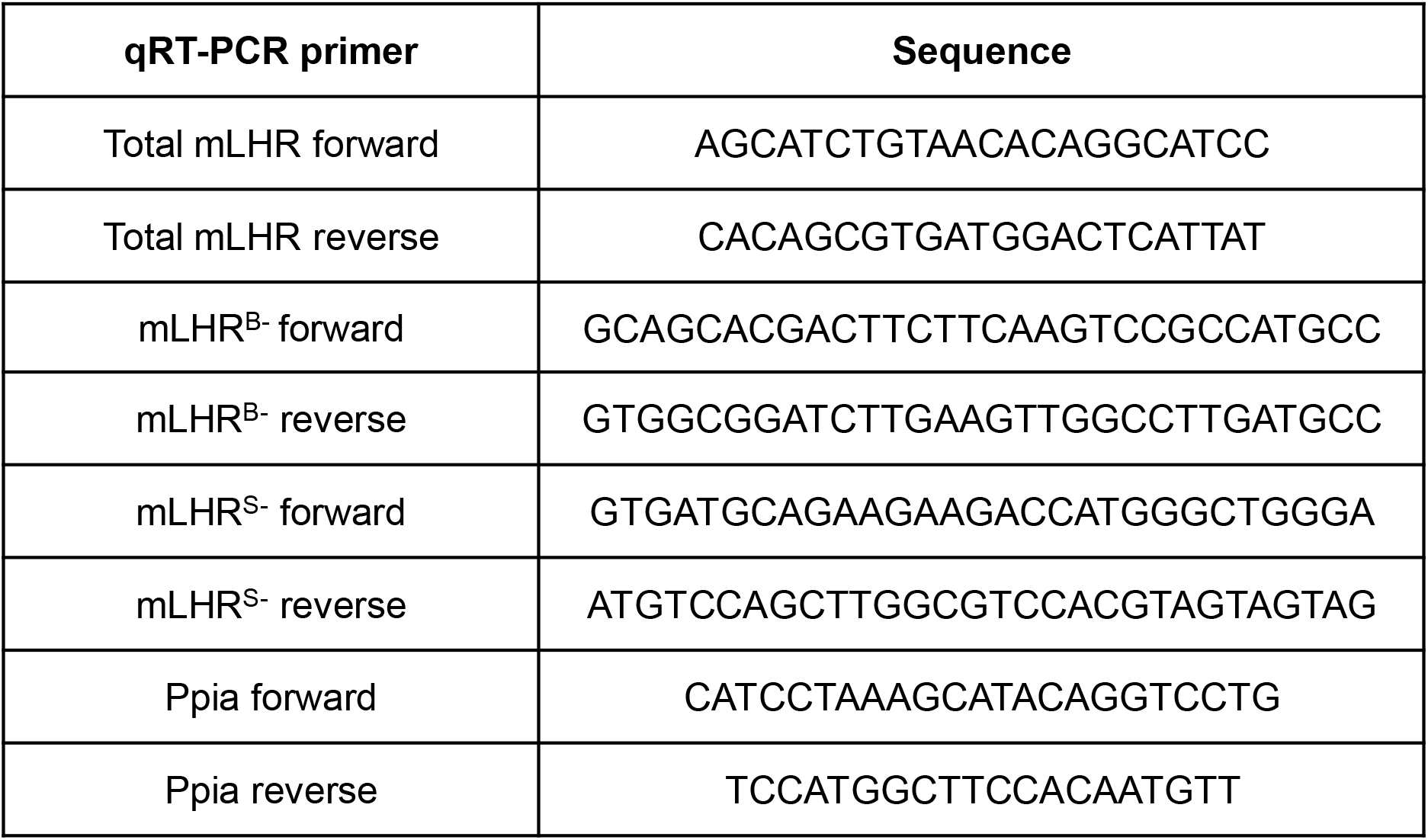
qPCR primers utilised for assessing WT mLHR, LHR^B−^ and LHR^S−^ expression levels.

### Statistical analysis

Statistical analyses were performed in GraphPad Prism, V8 using one-way ANOVA with Dunnett’s multiple comparison post-hoc testing. Data is represented as mean −/+ SEM for n= 4-8 animals. Statistical significance was determined as a *P* < 0.05.

## RESULTS AND DISCUSSION

### *In vivo* LHR homomerization via functional complementation in female mice

There is increasing evidence that GPCR di/oligomerization provides an important means to diversify/bias receptor functions^18,19^. Our previous study showed that ‘forced’ LHR homomerization via functional complementation was sufficient to restore the Leydig cell testosterone production and fertility of LuRKO male mice. However, whether LHR homomerization could restore the ovarian functions and fertility of female LuRKO remained unknown. Here, using the same BACs transgenic approach as previously employed, and functional complementation of binding (LHR^B−^) and signaling deficient (LHR^S−^) mutant LHRs (Figure 1A) we analyzed the effect of LHR^S−^ and LHR^B−^ co-expression in the LuRKO background on key reproductive parameters to assess this question.

**Figure 1.**
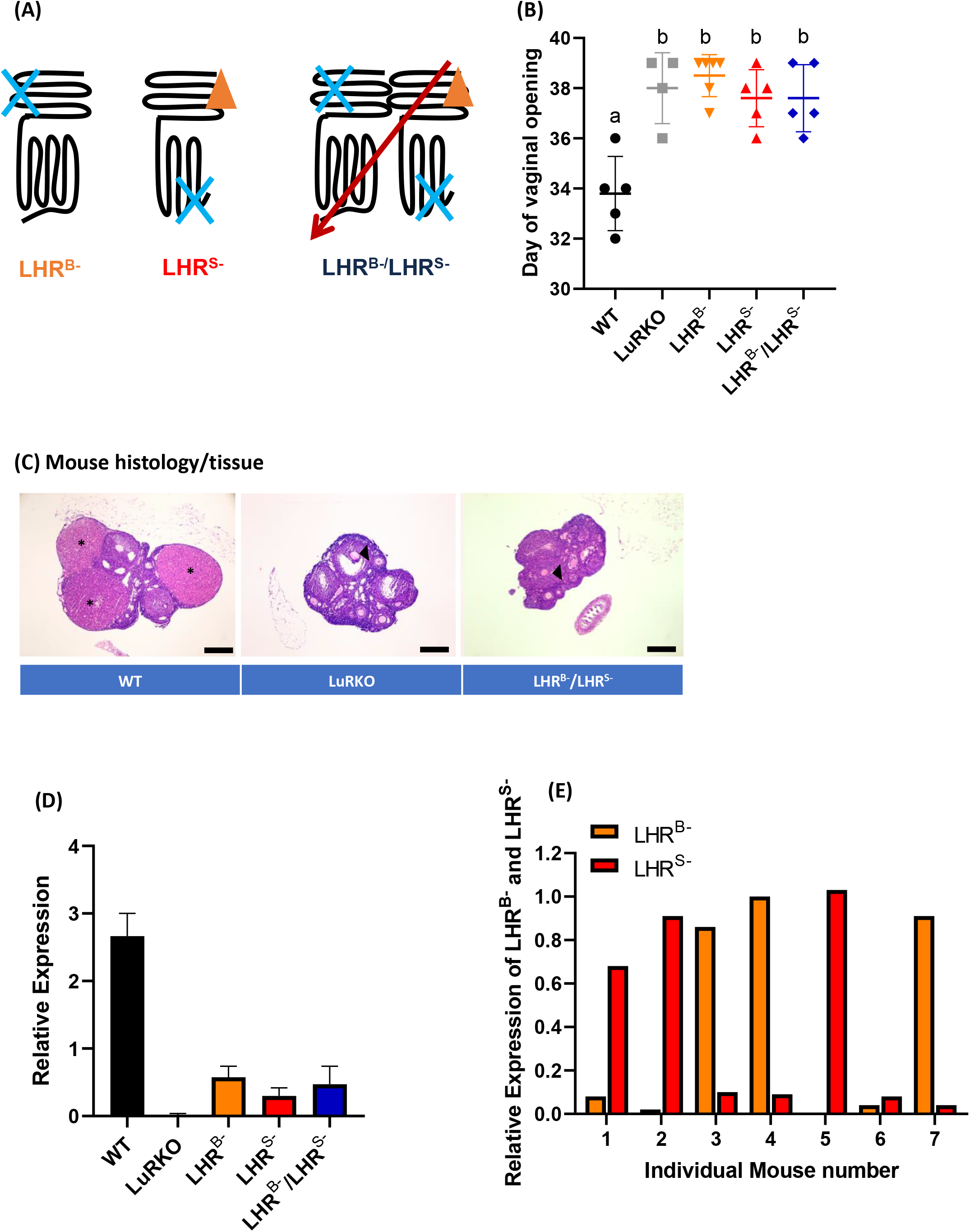
LHR functional complementation has no effect on the hypogonadal phenotype of female LuRKO animals due to low LHR^B−^/LHR^S−^ expression. **(A)** Schematic detailing LHR functional complementation and forced homomerization using LHR^B−^ and LHR^S−^ mutant receptors. The effects of LHR^B−^ and LHR^S−^ BACs co-expressed in a LuRKO background on **(B)** day of vaginal opening **(C)** Representative ovarian histology in 3-month-old control WT, LuRKO alone, or a LuRKO background expressing LHR^B−^/LHR^S−^ female littermates. * denotes presence of corpora lutea, arrows indicate failure of cumulus-oocyte-complex expansion **(D)** Relative LHR transcript levels and **(E)** relative transcript levels of LHR^B−^ and LHR^S−^ in single female mice co-expressing LHR^B−^/LHR^S−^. n=4-6 animals per experimental group, with statistical significance determined by One-way ANOVA with Dunnett’s multiple comparisons to the control and between groups. Alphabetic denotation a versus b= *** WT versus LuRKO animals, with no significant difference between LuRKO, LHR^B−^, LHR^S−^ and LHR^B−^/LHR^S−^ groups.

Because the postnatal sexual maturation is impaired in LuRKO animals, we first examined if LHR^B−^/LHR^S−^ co-expression could alter the timing of puberty via monitoring the day of vaginal opening. In wild-type and heterozygous LuRKO control animals, the first day of vaginal opening occurred at 33.8 ± 0.7 days (Figure 1B). In contrast, in LuRKO/LHR^B−^/LHR^S−^ females, the onset of puberty was delayed to 37.6 ± 0.6 days, mirroring the pubertal delay of 38.0 ± 0.7 days, observed in the LuRKO animals (Figure 1B). Single expression of either LHR^B−^ or LHR^S−^ in the LuRKO background, also had no effect on the pubertal delay observed in the LuRKO animals, suggesting that functional complementation failed to rescue the pubertal delay observed in female LuRKO animals.

To determine if co-expression of LHR^B−^/LHR^S−^ could rescue the anovulatory phenotype of the adult LuRKO female mice, ovaries from 3-4-month-old female mice were serially sectioned and histologically analyzed. The ovaries from the control heterozygous LuRKO and wild-type animals showed follicles of all stages of folliculogenesis and the presence of several corpora lutea (Figure 1C), indicating that ovulation had occurred. In contrast, in the LuRKO females co-expressing the LHR^B−^/LHR^S−^, although primordial, pre-antral and antral follicles were present, folliculogenesis was arrested at the large antral pre-ovulatory follicle stage of development (Figure 1C). Moreover, serial sectioning of the ovaries failed to locate the presence of any corpora lutea, indicating that these large antral follicles failed to undergo ovulation, as also observed in the LuRKO animals (Figure 1C).

The regulation of LHR expression is much more dynamic in the ovary than testis, requiring induced expression in granulosa cells in the mature large antral follicle^10,20,21^, the activation of multiple G protein-dependent pathways^22,23^ and transactivation of the EGFR^24–26^ for initiating key ovulatory pathway networks. We therefore wanted to interrogate if the expression levels of LHR^B−^ and LHR^S−^ were comparable to the wild-type (WT) LHR and whether this could account for disparity between the functional rescue observed between the male versus female mice. qPCR analysis revealed that the combined expression levels of LHR^B−^ and LHR^S−^ in the LuRKO/LHR^B−^/LHR^S−^ animals were just 20% of the wild type LHR expression levels (Figure 1D). The expression levels of LHR^B−^ and LHR^S−^ were therefore most likely insufficient to mediate the functional rescue, possibly due to lack of intact mechanisms to trigger LHR upregulation. Additionally, granulosa cell expression is required for coupling to Gαq and facilitating FSHR cross talk/heteromerization and EGFR transactivation necessary for progression of ovulation and high expression levels of LHR have been shown to be necessary for Gαq coupling^27^. Additionally, our previous in vitro data suggests that although Gαs activation is intact in WT and LHR^B−^/LHR^S−^ co-expressing conditions, LH (but not hCG)-dependent Gαq coupling is diminished in the latter^14^. Therefore, the lack of ovulation observed in LuRKO mice co-expressing LHR^B−^/LHR^S−^ may also reflect an impaired ability to activate LH-dependent Gαq signaling.

Our previous results have shown that the ratio of LHR^B−^ to LHR^S−^ directs the amplitude of Gαs and Gαq signaling observed^14^. We therefore determined the levels of LHR^B−^ and LHR^S−^ in individual mice co-expressing LHR^B−^/LHR^S−^. Surprisingly, we saw a wide range of expression levels, ranging from 1:22 LHR^B−^:LHR^S−^ to 45:1 LHR^B−^: LHR^S−^ (Figure 1E). Yet, in all cases the expression of individual or combined *Lhr* expression (by qPCR) was only a fraction of that of the WT in control animals (<25%). Our previous reported *in vitro* analysis of the ratiometric effects of LHR^B−^:LHR^S−^ expression suggested that an excess of cell surface LHR^S−^ :LHR^B−^ promoted more effective Gαs and Gαq signaling, contrasting with this *in vivo* data. However, the combined cell surface expression levels of LHR^B−^ and LHR^S−^ in cell lines analyzed were comparable to WT LHR, which may account for the disparity between our previous *in vitro* findings, and this study.

Overall, physiologically, these data suggest a much simpler male regulation of LHR functions, with a small amount of LHR signaling (< 1% receptor occupancy) sufficient to trigger testosterone generation, in concordance with previous studies^28^ and therefore rescuing spermatogenesis, as compared to the cyclical changes and more complex LHR actions evoked by higher LH levels and receptor occupancy in females. Additionally, whilst cellular compartmentalized expression of LHR and the FSHR occurs in males, in females LHR and FSHR are co-expressed in antral follicles of granulosa cells, which is hypothesized to be essential for the latter stages of ovarian follicle maturation and ovulation, via LHR-Gαq activation^23^. If such multifaceted LHR-dependent signaling activities are required in females (vs males) is a question that remains unanswered. However, with the advent of CRISPR, in all its forms, and thus the technologies to gene edit and regulate gene expression, it is conceivable to envision such experiments and again repurpose the LuRKO mice.

### LH ligands act explicitly via LHR

Recent studies have suggested that hCG may mediate its effects via alternative receptors to the LHR^29–31^. We therefore utilized our previously described hCGβ overexpressing transgenic mouse line, with circulating bioactive heterodimeric hCG concentrations approximately 40-fold higher than endogenous LH levels typically found in wild-type animals^15^. These mice were crossed into the LuRKO mouse background to determine whether high LH/hCG bioactivity could have any effects on the phenotype of the animals devoid of functional LHR. Assessment of postnatal sexual development showed a delay in the first day of vaginal opening of the LuRKO/hCGβ mice (Figure 2A) in comparison to WT (39.2 ± 2.1 days LuRKO/hCGβ versus 31.6±1.5 days WT, P<0.0001), mimicking the delay observed in LuRKO animals, rather than accelerated onset and precocious puberty usually observed with the hCGβ mice (^15^ and Figure 2A). Analysis of the uterine weights of LuRKO/hCGβ mice revealed a trend for decreased uterine weights in comparison to control and hCGβ littermates (Figure 2B), suggestive of estrogen deficiency in the LuRKO/hCGβ animals. Histological analysis of ovarian tissue revealed that the LuRKO/hCGβ ovaries contained follicles that were halted at the large antral follicle stage, with absent corpora lutea, an indicator of failure of ovulation, in concordance with the LuRKO animal ovarian phenotype (Figure 2C). Moreover, daily monitoring of the estrous cycle revealed that the LuRKO/hCGβ were acyclic (Figure 2D), mimicking the acyclicity and halted folliculogenesis of the LuRKO animals.

**Figure 2.**
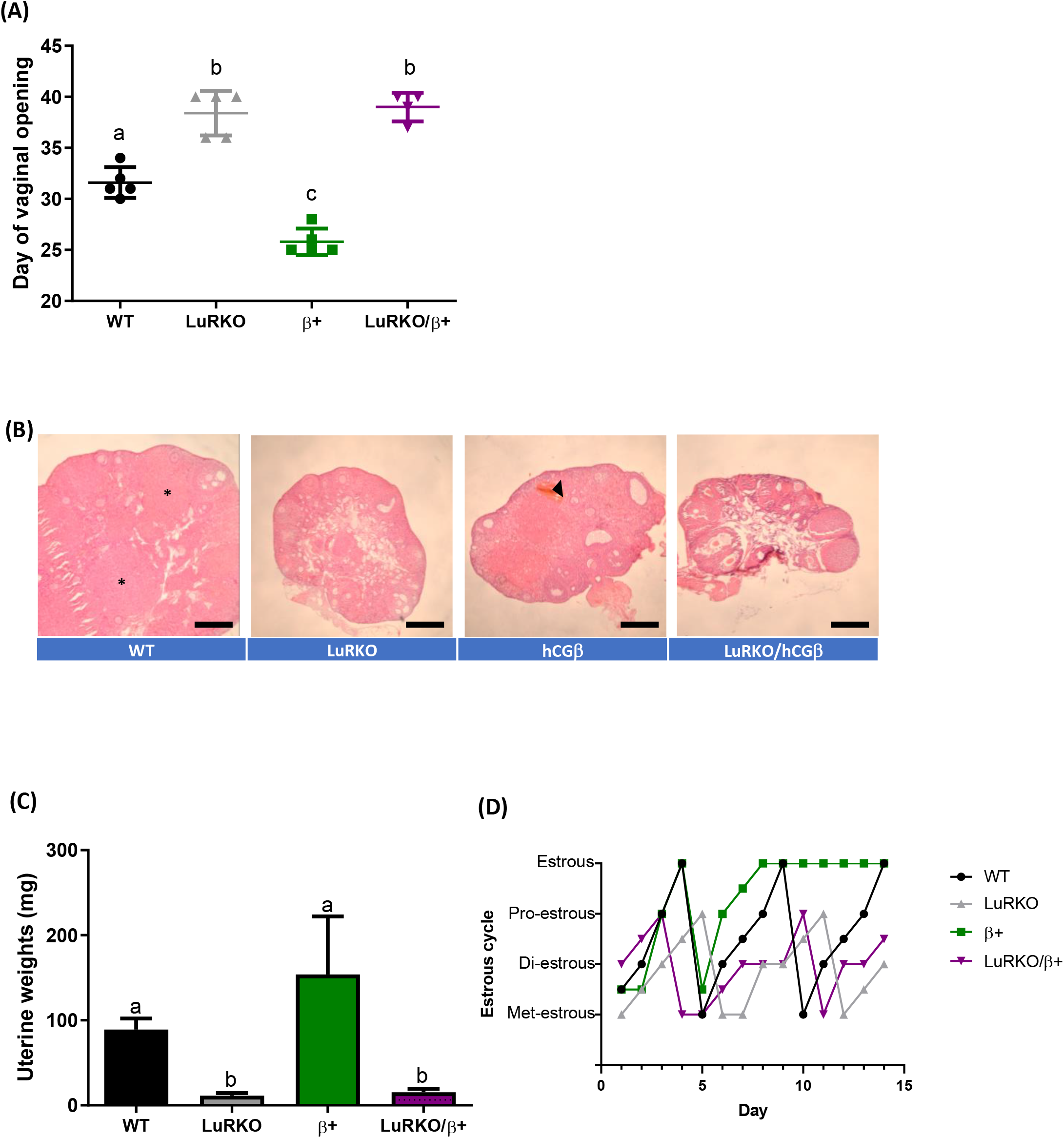
Enhanced ‘LH-like’ activity via hCG fails to rescue the hypogonadal phenotype of female LuRKO mice. The effects of hCG overexpression in the absence of LHR on key reproductive parameters as assessed by **(A)** Day of vaginal opening checked daily from d21 weaning and **(B)** Uterine weights in WT, LuRKO, hCGβ (β) and LuRKO/ hCGβ animals (LuRKO/β) females. **(C**) Histological characterization of ovarian sections. The ovaries of LuRKO/hCGβ mice were comparable to LuRKO animals, with follicles arrested at the large antral phase and lack of corpora lutea (* for corpora lutea in wild type animals). For the hCGβ mice, a hemorrhagic cyst, typical for mice with hCGβ expression, was also observed (arrowhead). Scale bars= 500 μm. WT included *Lhr^+/−^* females. **(D)** Representative examples of estrus cycles of each experimental group. Staging was Met= Metestrus; Di=Diestrus; Pro=Proestrus; Estr=Estrus. In case the phase has been difficult to define, it was marked between the two phases. n=4-5 animals per experimental group, with statistical significance determined by One-way ANOVA with Dunnett’s multiple comparisons to the control and between groups. Differential letter denotation equaled statistical significance between experimental groups, with a vs b= ****, a vs c= ***, and b vs c= ****.

These data provide no phenotypic evidence to suggest that high LH/hCG bioactivity can bypass LHR activation in animals devoid of functional LHR. Whilst our data using these mouse models cannot fully address whether in humans alternative functions for hCG exists, e.g. mannose-6-phosphate receptor-dependent activation in uterine natural killer cells cells^29^ or additional extragonadal roles proposed for hCG (reviewed in ^32^), it supports that the largest and most important actions of LH/hCG bioactivity are via the gonadal LHR, with most other effects observed downstream of LHR binding and signaling pathway activation. This is further supported by human inactivating or activating LHR mutations, where all the phenotypical abnormalities are connected to alterations in gonadal function^33–38^. Because LHR inactivation in the LuRKO mice is universal, our studies were not able to address the question about the functional significance of the extragonadal LHR expression (recently reviewed in ^39^).

### Constitutively activating mutations (CAM) of FSHR partially rescue the LuRKO female phenotype but not fertility

Our recent study has shown that the expression of CAM FSHR could rescue the hypogonadal phenotype and infertility of the male LuRKO mice^17^, suggesting a role for robust FSHR activity in compensating for missing LHR- and testosterone-dependent gonadal functions, including spermatogenesis. To address whether functional rescue could also take place in the female LuRKO/CAM FSHR mice, we assessed their key reproductive parameters. Analysis of the day of vaginal opening showed the LuRKO/CAM FSHR mice exhibited more variation in maturation timing, but it was not significantly different from WT females (33.5 ± 6.4 days vs 26.6 ± 2.2 days, respectively), while the delay was considerable in LuRKO female litter mates (45.4 ± 14.3 days, p < 0.05 between LuRKO and LuRKO/CAM FSHR mice) (Figure 3A). In addition, augmented uterine weights in the LuRKO/CAM FSHR females was observed in comparison to the LuRKO animals (Figure 3B), suggesting an enhancement in estrogen production in the former animals. Evidence of increased estradiol action due to CAM FSHR expression was also observed in mammary glands, which showed more advanced branching in LuRKO/CAM FSHR than in LuRKO females (Figure 3C, the upper row). In agreement with the previous data, histological analysis of ovarian tissue showed a rise in the number large antral follicles in the LuRKO/CAM FSHR females, in comparison to LuRKO animals (Figure 3C, the lower row). Clear separation of the cumulus-oocyte-complex was also observed (Figure 3C, the lower row, an arrow in an insert), indicating the potential further progression of these follicles in the presence of CAM FSHR, and enhanced estrogen production as a result of this. However, no corpora lutea were detected in LuRKO/CAM FSHR ovaries in contrast to CAM FSHR females that had often luteinizing follicles with trapped oocytes (Figure 3C and reference 15). Together the mouse models strongly suggest that intact LHR is required to mediate ovulation and excessive and constant FSHR action cannot replace it. As a result, the LuRKO/CAM FSHR females were acyclic (Figure 3D), mirroring the arrested cyclicity observed in LuRKO female mice. As expected, the LuRKO/CAM FSHR females set for breeding with fertile males failed to present with vaginal plugs, indicative of the lack of copulatory behavior in the females, nor pregnancy suggesting the females were infertile (data not shown).

**Figure 3.**
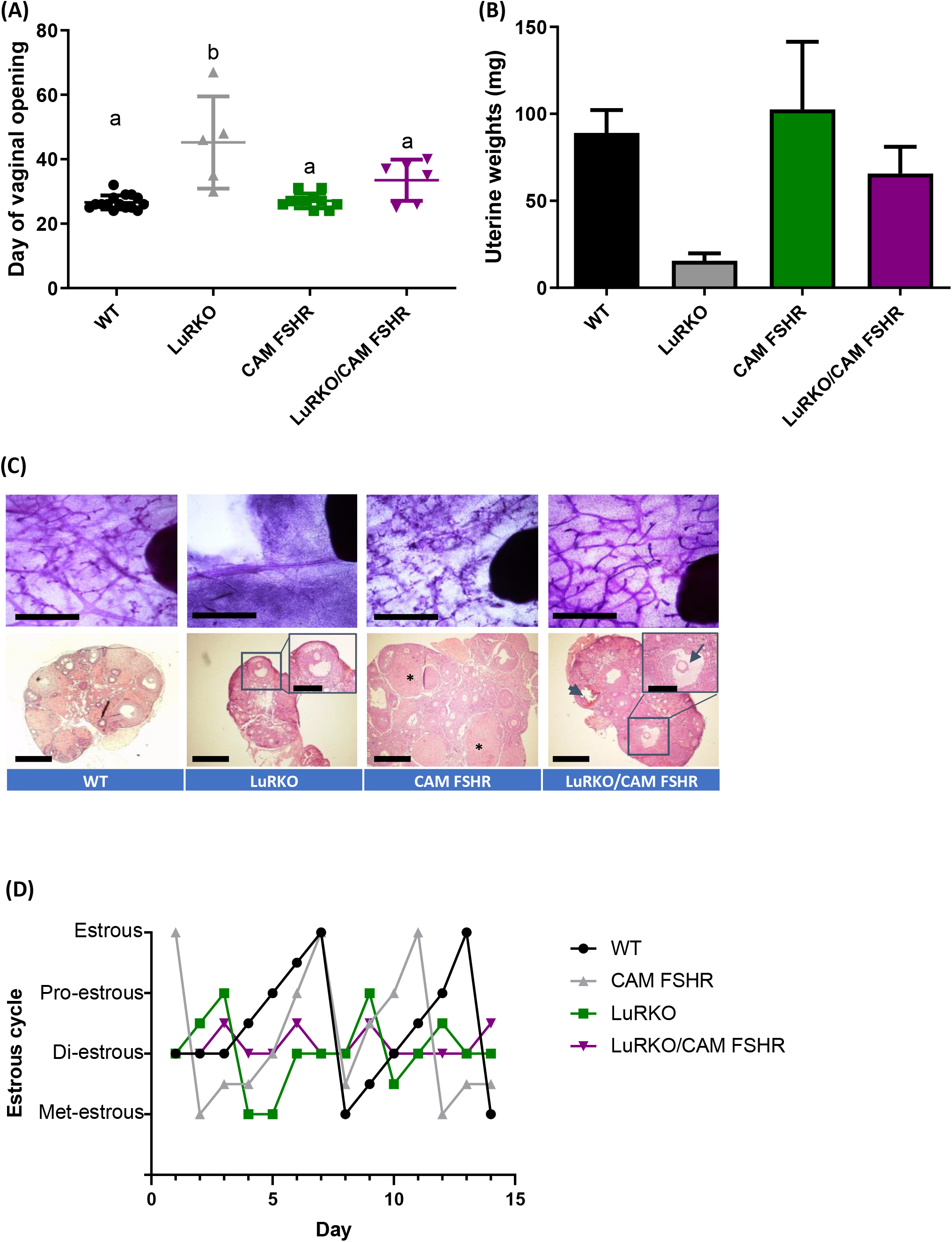
Constitutive activation of FSHR in the absence of LHR partially restores ovarian function and mammary gland development. Key reproductive parameters were assessed via **(A)** Day of vaginal opening and **(B)** Uterine weights in WT, LuRKO /CAM FSHR, CAM FSHR, and LuRKO females. **(C**) Histological characterization of mammary gland and ovaries of WT, LuRKO, CAM FSHR, LuRKO/CAM FSHR and mice. Upper row: Whole mount in situ staining of the mammary glands demonstrates the rescue of branching in LuRKO/CAM FSHR female mice, versus rudimentary development in LuRKO females. Lower row: The ovaries of CAM FSHR mice are typically larger than those of WT mice and contain multiple developing follicles and luteinizing follicles (*). The ovaries of LuRKO/CAM FSHR mice have higher number and more advanced follicles (an arrow in an insert) than LuRKO mice, but lack corpora lutea. A hemorrhagic cyst, typical for mice with CAM FSHR, is also seen (arrowhead). Scale bars: the upper row 1 mm; lower row 500 μm and inserts 250 μm. WT included *Lhr^+/−^* females and CAM FSHR in WT and *Lhr^+/−^* backgrounds. **(D)** Representative examples of estrus cycles. Met, Metestrus; Di, Diestrus; Pro, Proestrus; Estr, Estrus. Where the phase has been difficult to define, it has marked to be between the two phases. CAM FSHR females demonstrated variable cycles from semi-normal to acyclic, in line with our previous publication^16^, of which one has been shown here. n=3-15 animals per experimental group, with statistical significance determined by One-way ANOVA with Dunnett’s multiple comparisons to the control and between groups. Differential letter denotation equaled statistical significance between experimental groups.

The enhanced estrogenic activity as evidenced by increased uterine weights and progression of follicular maturation suggests that constitutive activity of FSHR advanced follicle maturation in the absence of LHR. This boosted follicular maturation is in line with our previous work showing accelerated follicle maturation in the CAM FSHR mice^16^ and provides additional insight into how constitutive activation of FSHR can partially fulfil LHR-dependent roles within antral follicle maturation to the pre/ovulatory follicle. However, intact LHR/LH signaling is ultimately required for ovulation. As previously discussed, the physiological roles of LHR/FSHR heteromers has been postulated, with proposed roles in modulating signal specificity within the ovulatory follicle^40–42^. Although enhanced ligand-independent cAMP signaling has been previously described for this CAM FSHR^16^, the constitutive activation of additional FSH/FSHR pathways such as PI3 kinase/AKT, β-arrestin and ERK-MAPK, and Gαq mediated Ca^2+^ and PKC signaling remains unknown.

Ovarian function of the CAM FSHR/LuRKO mice agrees with our previous findings on FSH-treated LuRKO mice^43^, where FSH stimulation was unable to promote follicular maturation beyond the antral stage. This contrasts with previous studies with hypophysectomized rats and mice, where ovulation and luteinization could be induced by treatment with recombinant FSH without LH^44–46^. This prompts the necessity of intact LHR expression, even without ligand, for ovulation. One possibility is that this response requires LHR/FSHR heterodimerization to transduce the complete FSH signal.

## CONCLUSIONS

This study has utilized three approaches to interrogate the mode and nature of LHR signaling in modulating ovarian function. It suggests the importance of the spatial-temporal changes in LHR expression and the requirement for LH-dependent pleiotropic signaling for mediating ovulation. Yet, outstanding questions remain surrounding the functional relevance of LHR homomerization and LHR/FSHR heterodimerization within the ovary, a question that will no doubt be answered by the rapidly evolving gene editing approaches that are now available. Additionally, although LHR is essential for ovarian function, the postulated extragonadal roles of gonadotropin hormones in pregnancy and relevance of gonadotropin hormone receptors for mediating these roles remains to be determined.

## ACKNOWLEDGEMENTS

We gratefully acknowledge BBSRC grant number BB/1008004/1 and Wellcome Trust grant number 082101/Z/07/Z for funding these studies.

## Notes

**Funding**: BBSRC grant number BB/1008004/1 and Wellcome Trust grant number 082101/Z/07/Z to IH.

### Competing Interest Statement

The authors have declared no competing interest.

